# Systematic comparative benchmarking of computational methods for the detection of transposable elements in long-read sequencing data

**DOI:** 10.1101/2025.09.29.679192

**Authors:** Nogayhan Seymen, Renato Santos, Ramya Lakshmanan, Simon Topp, Ammar Al-Chalabi, Ahmad Al Khleifat, Gerome Breen, Richard JB Dobson, John P Quinn, Mohammad M. Karimi, Alfredo Iacoangeli

## Abstract

**Background:** Mobile element insertions, particularly transposable elements (TEs) such as Alu, LINE-1 (L1), SVA, and endogenous retroviruses (ERVs), represent a major source of human genetic variation and have been implicated in evolution, genomic instability, and disease. Although long-read sequencing generally outperforms short-read sequencing for the characterisation of such elements, their accurate detection with long-reads remains challenging, with different computational tools adopting varying approaches and producing divergent call sets. As gold standards currently do not exist for TE detection, benchmarking these methods is essential to understand their strengths, limitations, and biases. Here, we systematically evaluate the performance of available state-of-the-art TE detection tools on both simulated and real human genome data using highly characterised samples from the Genome in a Bottle consortium, population level reference databases and an in-house collection for which matching short-read sequencing data are available.

**Results:** Our results show significant differences in calling strategies, leading to substantial variation in precision, recall, and the spectrum of TE families detected across tools. Our benchmark also displays the differences between short-read and long-read calls, highlighting the importance of appropriate method selection.

**Conclusions:** The benchmarking results presented here will aid TE researchers make better informed decisions on which tool to use in their long-read TE analyses. Strengths and limitations of different tools have been highlighted in depth as well as their computational requirements, which will result in less time spent finding the best tool for the job and promote faster TE research.

## Background

Transposable Elements (TEs) are mobile genomic sequences that are capable of self-replication either by cut-and-paste (DNA transposons) or copy-paste (retrotransposons (RTEs)). It is estimated that nearly half of the human genome is composed of transposons, albeit less than 0.05% of them are believed to remain active[1]. There are four actively mobile and well-studied RT families: Alu, L1, SVA and ERV (Figure 1). Owing to their different structures, different families of RTEs vary greatly in length. While Alus are approximately 300bp, making them the shortest TE family, ERVs are around 10,000bp long, making them especially difficult to detect using short-read sequencing (SRS) techniques[2]. However, long-read sequencing (LRS) can generate reads ranging from 10kb to 1Mb long, which not only enables the accurate sequencing of complex regions in our genome, but also structural variations (SVs), including TEs[3].

**Figure 1:**
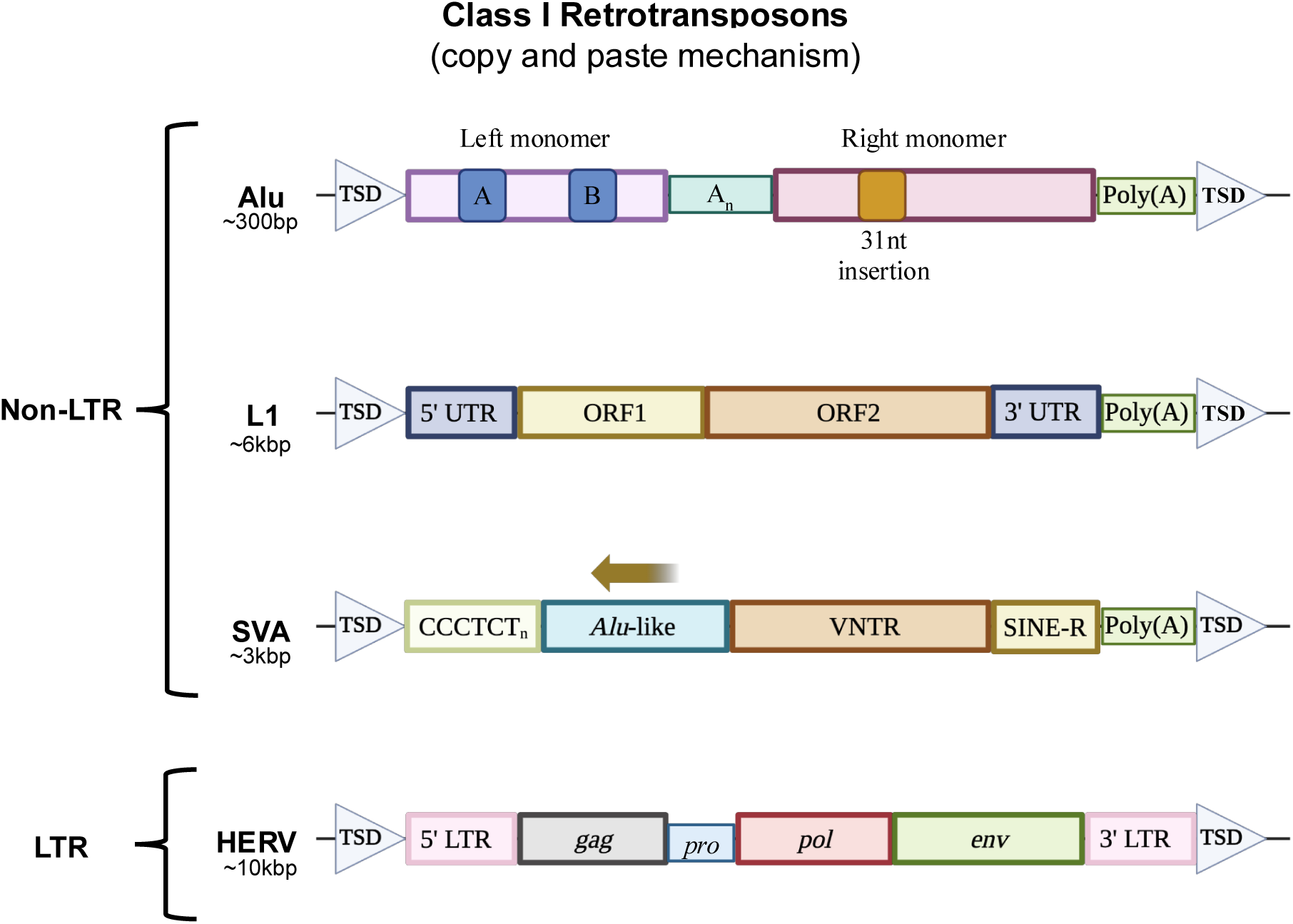
Class I Retrotransposon genomic structures.

**Figure 2:**
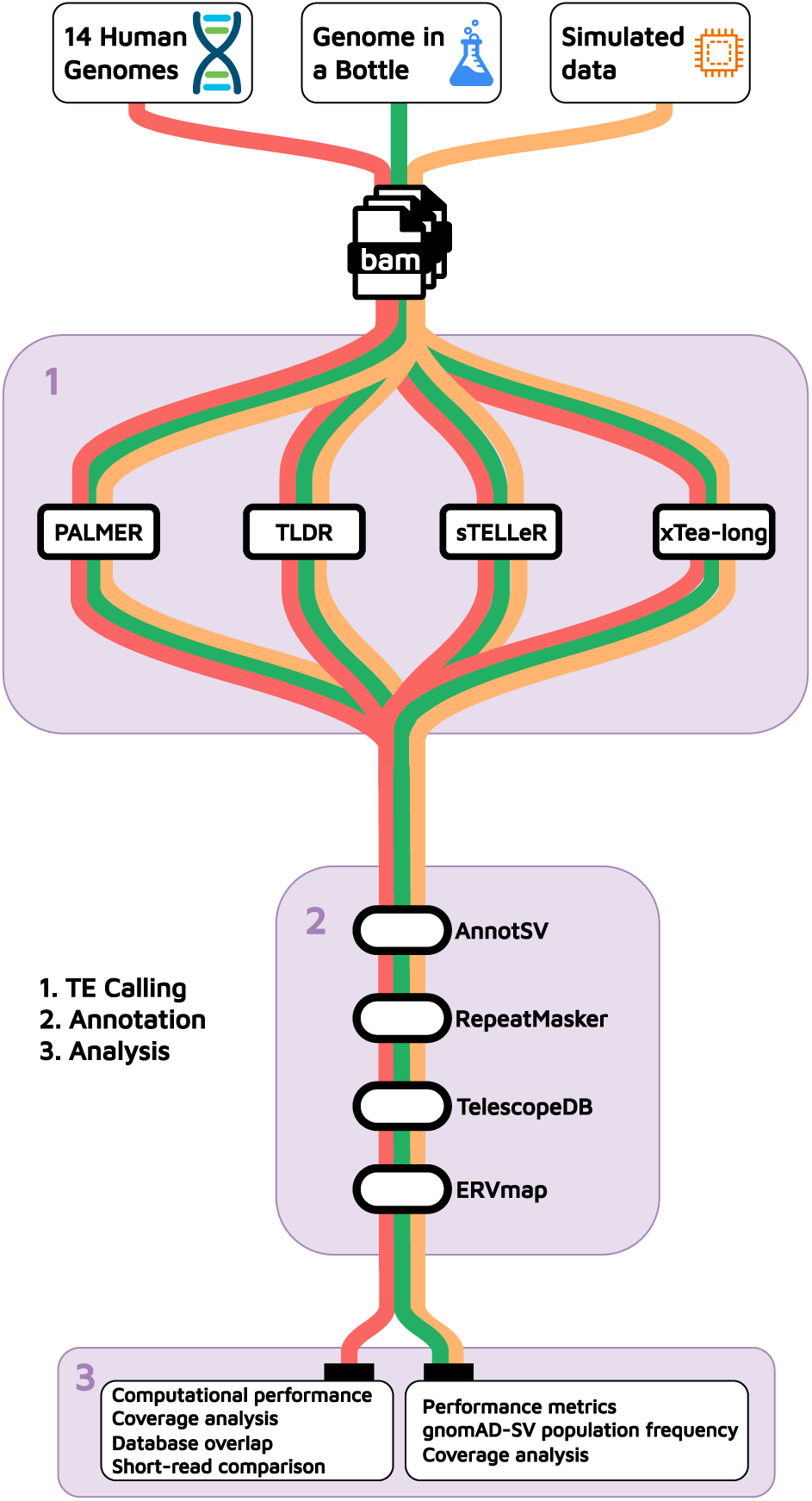
Overview of the workflow for this study.

Evidence exists involving TEs in numerous biological processes including human diseases. For example, L1 insertions into the *F8* gene were shown to cause haemophilia[4], and Alu insertion in exon 6 of the *NF1* gene is proven to cause neurofibramatosis[5]. ERVs are remnants of ancient retroviral infections and many are still capable of transcription via a number of mechanisms. For example, some ERVs can be activated by certain epigenetic factors [6], causing elevated levels of transcripts. An example of this was discovered by Phan *et al.* where elevated levels of the env transcript of a specific family of human ERV (HERV) called HERV-K, were found in the blood serum of Amyotrophic Lateral Sclerosis (ALS) patients[7]. These examples represent well how critical to accurately detect TEs is to better understand the mechanisms behind human diseases. However, apart from being present or absent in genomic regions, TEs can also be polymorphic. Both reference TEs, which are TEs annotated in the human genome assembly, and non-reference TEs, can contain polymorphisms or represent novel insertions unique to individuals. Detecting non-reference TEs is crucial for understanding genetic diversity, population structure, and disease-associated insertion events[8,9].

Long-read sequencing has gained momentum in the last decade, and individual base calling accuracy has become comparable to SRS[3]. This has led to an influx of new tools developed specifically for LRS applications including non-reference TE detection. Benchmarking of newly developed computational tools can be influenced by the selection of comparative methods, which may not fully represent the diversity of available approaches. As most long-read non-reference TE detection tools have only been introduced within the past year, comprehensive and systematic benchmarking studies remain lacking. We hereby aim to close this gap by performing a thorough evaluation of the available tools for non-reference TE detection using long-read sequencing data. We have run four specialist tools, PALMER[10], TLDR[11], xTea[12] and sTELLeR[13] with three different datasets: simulated long-read sequences, sequencing data of the HG002 sample from the Genome In A Bottle (GIAB) consortium[14] as well as an in-house collection of 14 samples from the London Neurodegenerative Diseases Brain Bank (LND BB)[15,16] sequenced with Oxford Nanopore Technologies (ONT). We assessed their performance using population-level variant frequencies from gnomAD-SV database and high confidence sets from the Genome in a Bottle consortium, as well as exploring the agreement across tools and from matching SRS samples in our in-house data.

## Results

### Benchmarking design

The purpose of this study was to benchmark available tools for non-reference TE detection that are specifically developed for long-read technologies. To that end, we have run all available tools, PALMER, TLDR, xTea, and sTELLeR on 3 different datasets and compared them based on computational efficiency, performance and overall ease-of-use. We are aware that as of performing this benchmark study, there are two more tools that fall into this category, LOCATE[17] and TrEMOLO[18], which were not included in this study. LOCATE is currently available as a preprint and undergoing revision and TrEMOLO could not be implemented into our workflow due to repetitive technical issues with running the tool. Both tools represent promising approaches and will be important to include in future benchmarking analyses should they improve user experience and documentation.

We used both simulated and real data for benchmarking. If two TEs were predicted to be within 50 bp from each other, they were considered the same TE, therefore, TEs predicted to be within 50 bp from a simulated TE or a high confidence call, depending on the experiment, were considered true positives (TP). We applied a relatively strict gap allowance to capture differences among tools while maintaining a narrow true positive (TP) window. To simulate the LRS data, we used a custom script to inject TEs into the reference genome which we then used to simulate reads, using pbsim3[19].

1409 Alu, 1266 L1, 645 SVA and 638 HERVs were inserted randomly into the GRCh38 genome this way. To observe if coverage effects the performance of the tools, we simulated LR samples with 5x, 10x, 20x and 40x coverage. Secondly, we downloaded the ultra-long ONT (∼63x mean coverage[20]) BAM files for the son (HG002) from GIAB. Finally, we used 14 ONT samples from LND BB to compare the tools in a “real-world” dataset.

### Computational performance and usability

We used all tools with default parameters except for number of threads and TE reference fasta where applicable. Among the 4 tools, only TLDR and xTea are capable of multithreading. xTea is the only tool that does not allow changing the TE reference fasta (at least not without changing source code) (Table 1). In order to remove the effect of the TE reference provided on the performance of the tools, we used the same TE reference for the other 3 tools.

**Table 1:**
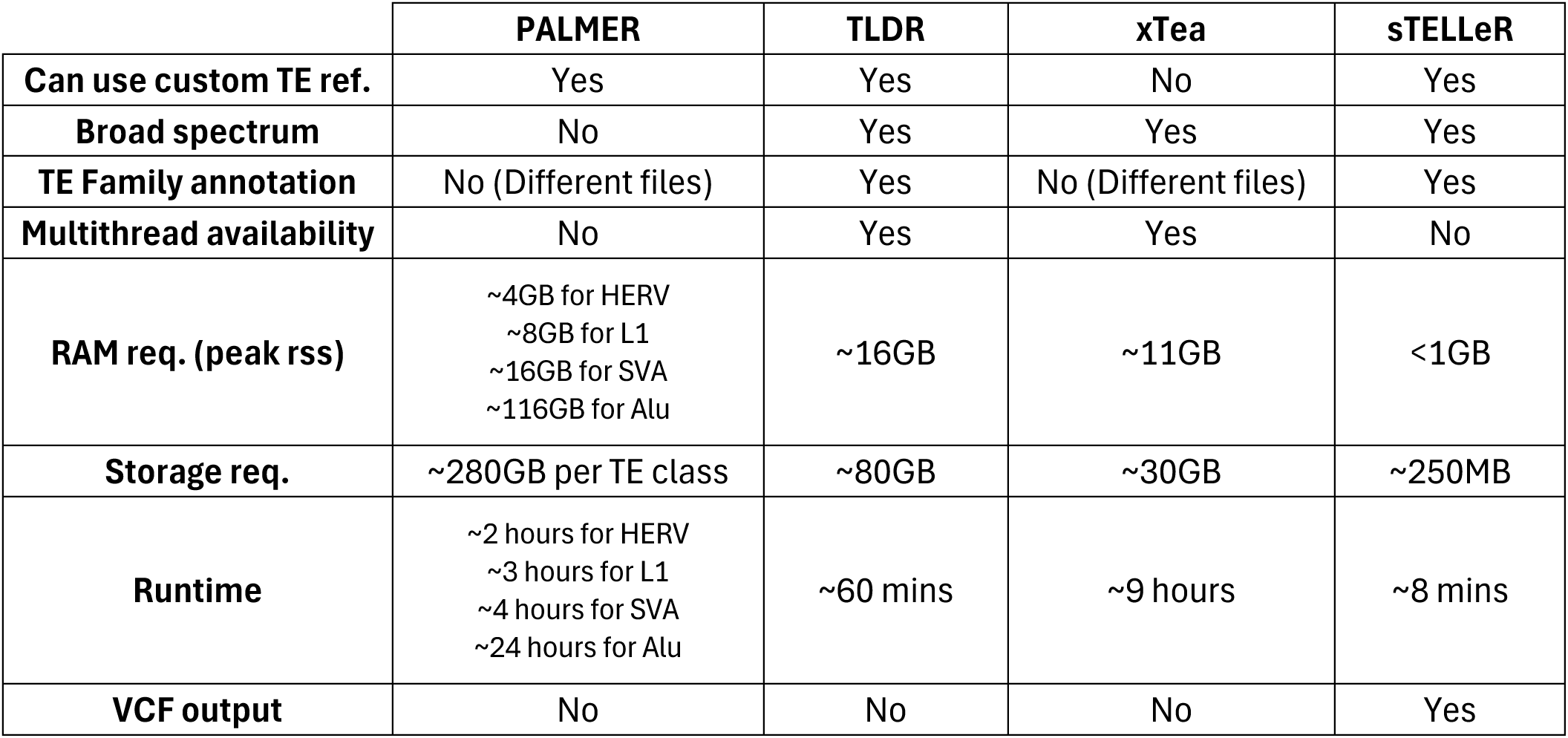
Computational performance and usability comparison table for PALMER, TLDR, xTea and sTELLeR. RAM req., Storage req. and Runtime values are rounded means for the 14 LND BB samples.

Regarding computational requirements and ease of use, using the 14 Brain Bank samples, PALMER is the most resource heavy tool, with more than 100 GB of Random Access Memory (RAM) required to run for Alu detection (Table 1). It also requires on average about 250 GB of hard drive space per TE class, which results in ∼1Tb per samples if run all four TE families. PALMER is also the only tool that needs to be run separately for each TE family. Therefore, the runtime and memory requirements vary from 3 GB of memory and 2 hours of runtime for HERVs to upwards of 100 GB of memory to about a day of runtime for Alus. sTELLeR is by far the least resource intensive tool, using just 1 thread, less than a GB of memory and takes on average less than 10 minutes to run. TLDR and xTea use moderate resources but there is a big difference in their runtimes, with TLDR taking less than an hour to complete while xTea takes a few hours (Table 1)(Figure3). Coverage and related to it input file size directly affect the computational requirements regardless of the tool used. As can be seen in Figure 3, in every measurement other than xTea’s disk read which does not seem to be affected by coverage, singleplex (higher coverage) samples have higher resource usage. One important difference between xTea and the other tools is that it does not allow for the use of a custom TE reference fasta file. As for ease of use, all 4 tools were relatively easy to set up and start using, with sTELLeR having an official docker container, and the other tools have appropriate installation guides in their GitHub pages as well as unofficial Singularity containers at quay.io (PALMER: ymostovoy/lr-palmer, xTea: ymostovoy/lr-xtea, TLDR: ymostovoy/lr-tldr) which we used for this benchmark. xTea requires an additional “setup” script to be run for each sample, which creates the required files and directories as well as the Slurm submission script required to run for all submitted samples. For downstream analysis, sTELLeR has a neat VCF output while other tools have custom tables as outputs, which may be an item to consider if there are intended downstream analysis steps.

**Figure 3:**
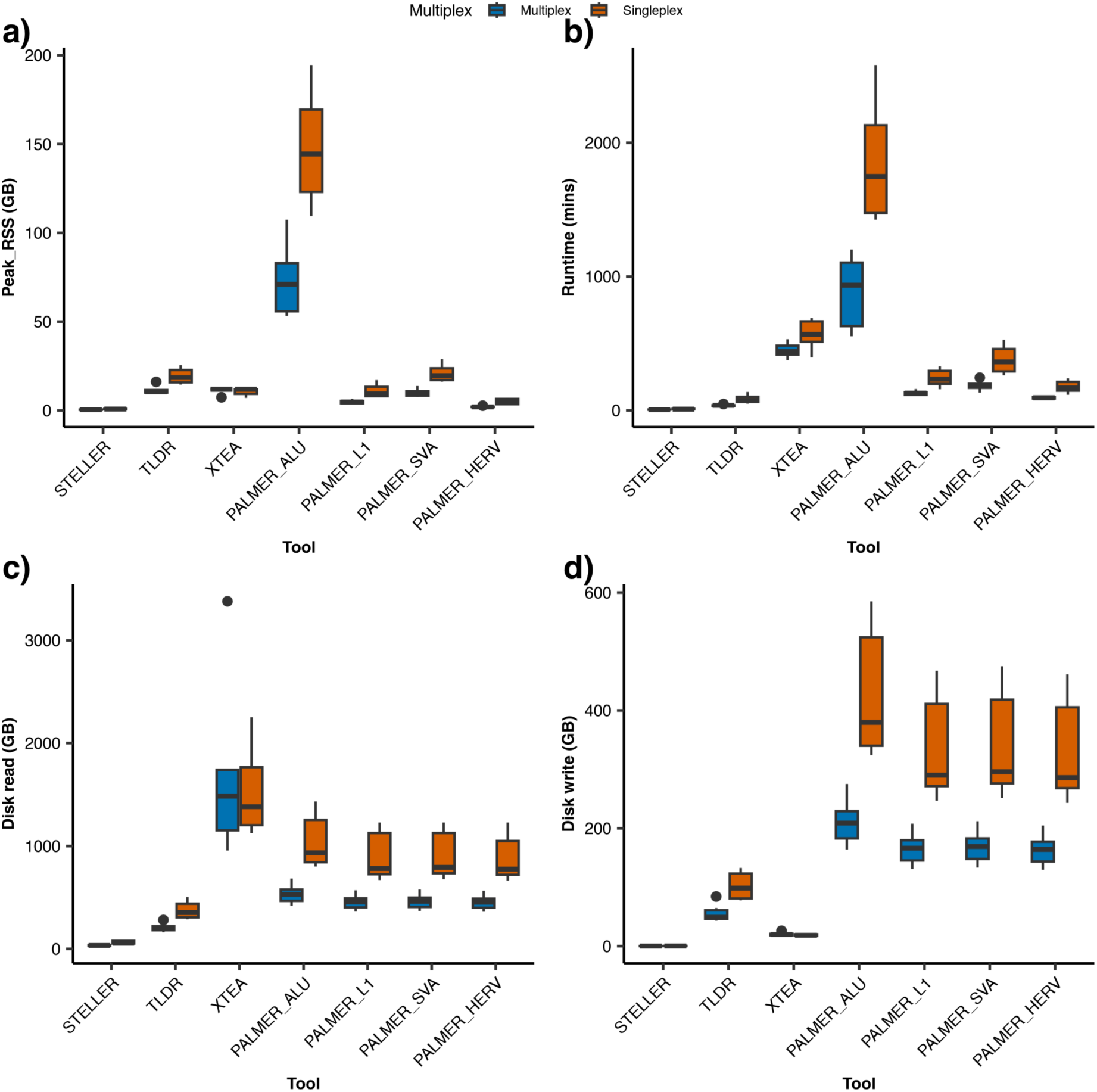
Computational performance of different tools for the 14 LND BB samples, split by multiplex status. a) Peak RSS RAM usage in GBs. b) Total runtime in minutes. c) GBs read from disk during running. d) GBs written to disk while running.

### Simulated data performance

All tools perform well with simulated data, though TLDR seems to be lagging a little bit behind (Fig. 3a). The F_1_-scores range from 96.3% - 97.8% for xTea, sTELLeR and PALMER while the highest F_1_-score for TLDR is 94.4% at 10x coverage (Supp. Table 1a). The performance overall increases with coverage, the only exception being the precision for PALMER which has a downward trend as coverage increases. xTea performed best with simulated data with 95.8% recall, 99.9% precision and 97.8% F_1_-score. Regarding individual TE families, HERV detection performance is highest in xTea, sTELLeR and PALMER with xTea having the top spot with 98.5% F_1_-score, 99.95% precision and 97.1% recall (Supp. Table 1b). It is worth mentioning that both xTea and sTELLeR perform best at 40x while PALMER has the highest score at 20x and TLDR at 10x which could result from their algorithmic differences but may indicate that while TLDR and PALMER have an optimal coverage window around the mentioned coverages, xTea and sTELLeR perform well with higher coverages. We also investigated the ensemble performances, combining the results of 4 tools in any combination, taking TEs detected in both/all tools as positive (Fig. 3b). PALMER + xTea combination performed best with 97.8% F_1_-score, 99.96% precision and 95.7% recall at 40x coverage, followed by 10x and 20x results of the same combination (Supp. Table 1c). Similarly, HERV detection is highest in all combinations of xTea, sTELLeR and PALMER, with PALMER + xTea being highest at 98.5%% F_1_-score, 99.95% precision and 97.1% recall (Supp. Table 1d).

### Genome in A Bottle data performance

For evaluating the performance of the tools on GIAB HG002 data, we performed three different analyses using three different truth sets. The first consisted of TEs from the “highest confidence” structural variant (SV) calls published by Zook et al[21]. To generate this set we annotated this confidence SV list using AnnotSV[22] for TE family annotation. The resulting SVs that had the same TE annotation in both “Repeat_type_left” and “Repeat_type_right” from AnnotSV were considered TE insertions and included in this truth set. To generate the second set, we filtered the same annotated SV calls to derive a higher confidence truth set by taking only calls that are called by 3 out of 4 tools and are in the GIAB SV list. As third set, we used high confidence L1 calls obtained using PALMER[23] and published on GIAB servers.

For the first truth set, there is similar performance throughout the 4 tools with sTELLeR having the highest F_1_-score at 44%. Overall, there is a tendency to capture more true positives at the expense of increased false positives which could be expected as we are comparing a non-filtered set of TE calls from the tools to an extensively filtered “high-confidence” SV call set. As a result, precision is generally low with sTELLeR and xTea sharing the highest precision here at 35%. On the contrary, there is better recall throughout with PALMER being highest with 71% and xTea at 50% (Figure 4a). These results highlight substantial trade-offs between recall and precision across tools, with no single method excelling equally in both metrics. To assess whether consensus calling could improve accuracy, we evaluated combinations of tools using the intersection of their call sets (Figure 4c). In this conservative approach, a TE call is considered true only if detected by all tools in the combination. This strategy generally improved precision compared to single-tool results (rising from ∼0.29–0.35 to ∼0.33– 0.36), reflecting a reduction, although still suboptimal, in false positives. However, the increased stringency came at the cost of reduced recall, as many true positives detected by only one tool were excluded. The highest recall (0.55) and balanced performance were observed for the PALMER + TLDR combination (F_1_-score = 0.43, Precision = 0.36), closely followed by PALMER + sTELLeR (Recall = 0.54). As more tools were added to the intersection (e.g., three or four tools), recall dropped further to ∼0.38–0.46, while precision gains were modest. This pattern highlights the trade-off between recall and precision inherent in strict consensus-based TE calling.

**Figure 4:**
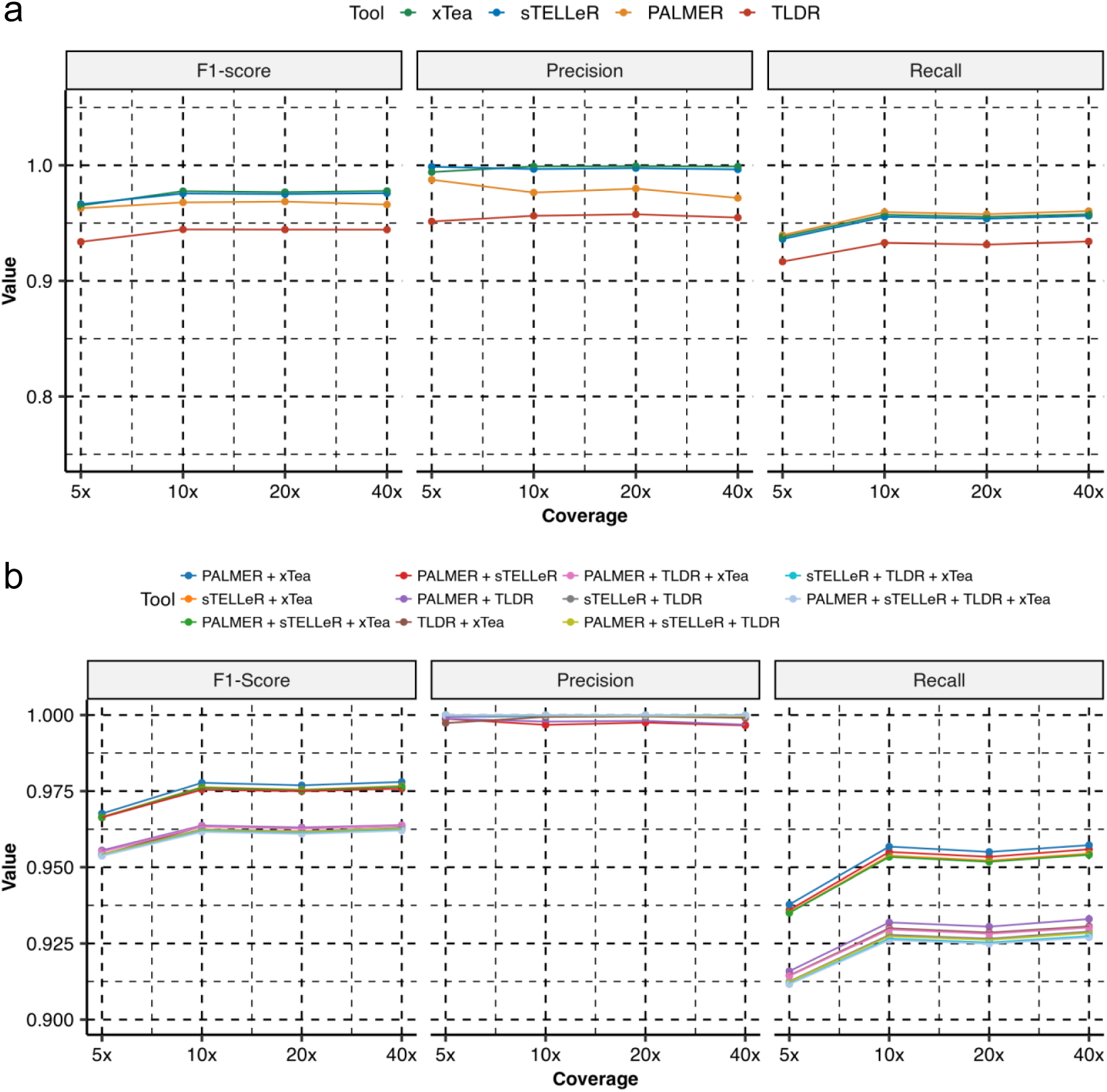
Performance metrics for simulated data. a. Precision, Recall and F1-scores for PALMER, TLDR, sTELLeR and xTea for 5x, 10x, 20x and 40x coverage. b. Precision, Recall and F1-scores for combinations of the four benchmarked tools for various coverages.

As expected, there is a dramatic increase in performance with the second approach where we filtered the same annotated SV call set to derive a stricter truth set. F_1_-scores increase to 85-95% with sTELLeR being 6% higher than its closest rival here. With a precision of 93% and recall of 96%, sTELLeR is the best tool when we want to validate a curated list of detected TEs. Two interesting points that follow the overall theme for the tools are 1) PALMER has the highest recall at 98%, which is expected as it is quite permissive in calling TEs; 2) xTea has the highest precision at 94% albeit a significantly reduced recall compared to the other tools, highlighting xTea’s strict calling algorithm (Figure 5).

**Figure 5:**
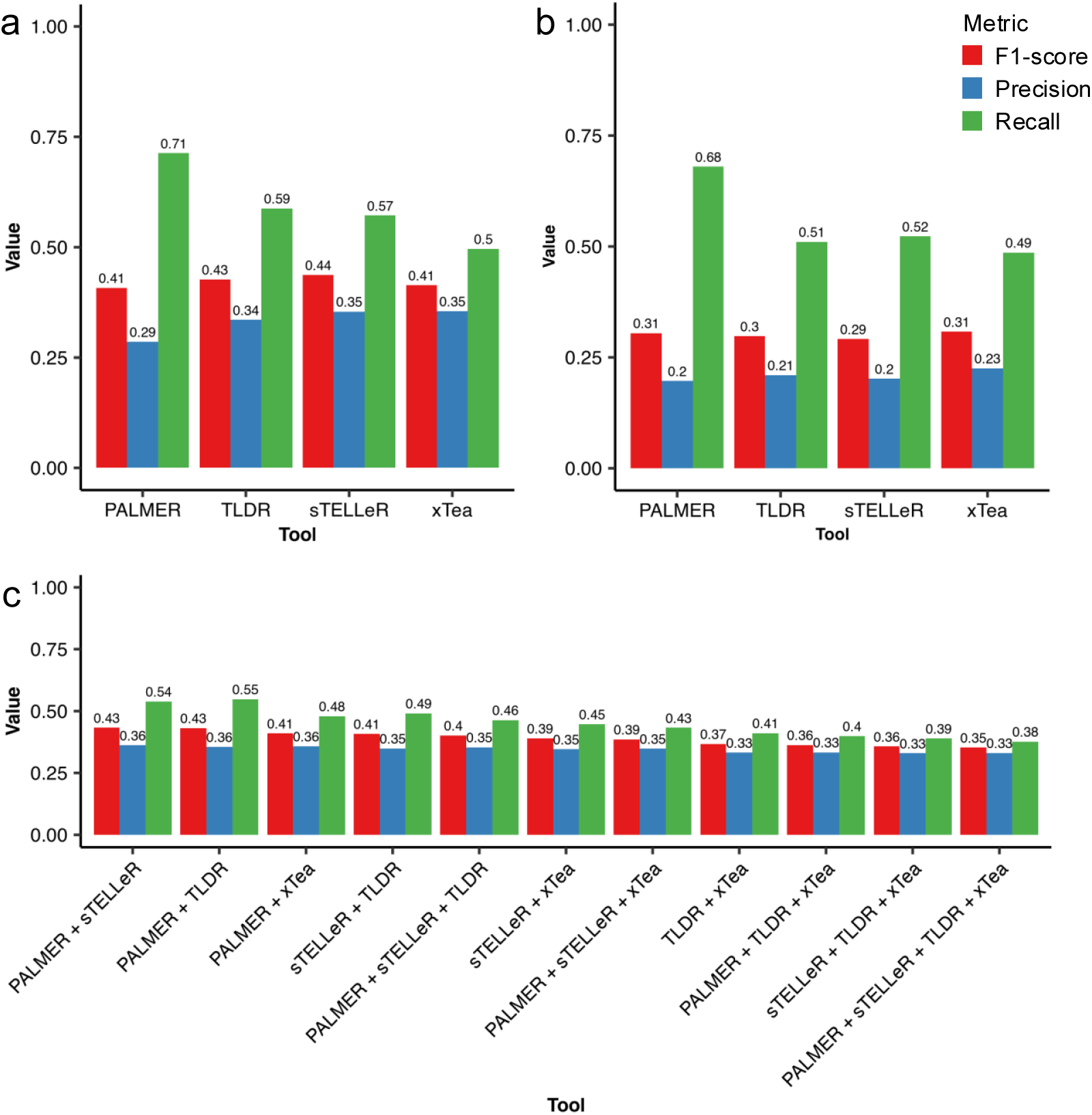
Performance metrics for GIAB data. a. Precision, Recall and F1-scores for PALMER, TLDR, sTELLeR and xTea for the GIAB data using annotated high-confidence SV calls as truth set. b. Precision, Recall and F1-scores for PALMER, TLDR, sTELLeR and xTea for the GIAB data using curated PALMER L1 list as truth set. c. Precision, Recall and F1-scores for combinations of the four benchmarked tools for the GIAB data using annotated high-confidence SV calls as truth set.

When we inspect separate TE families, Alu performance tends to be better for all tools with the exception of sTELLeR having the same F_1_-score in both Alu and SVA detection at 95%. In line with the rest of the benchmark, PALMER has the lowest precision which is more pronounced in L1 with the lowest precision of 63% (Supplementary Figure 1). While Alu, L1 and SVA performances are higher in sTELLeR, peak performance in HERVs comes from xTea with a very balanced 93% F_1_-score, 94% precision and 92% recall, once again exhibiting the strengths and weaknesses of the different algorithms. We’ve then assessed the performance in consensus calling as before, where PALMER + sTELLeR performed the best (F_1_-score = 0.95, Precision = 0.96, Recall = 0.94) (Supplementary Figure 2). With the exception of TLDR + xTea – the two most stringent callers in the benchmark, we can see that whenever more than 2 callers are combined, precision increases to 100%, which could indicate to a preferable strategy if the aim is to generate a highly reliable, high-confidence call set with minimal false discovery, accepting the corresponding reduction in recall.

### London Neurodegenerative Diseases Brain Bank data evaluation

We used 14 ONT (6 multiplex + 8 singleplex) samples from LND BB to run the tools on and compare the called TE sets. Mean depth for multiplexed samples was 9.76x while singleplex samples had approximately twice the coverage at a mean of 19.68x[24]. More detailed information about the sequencing and quality of these 14 samples can be found in the original paper that describes the dataset [24].

First, we checked if depth of coverage of samples was correlated with the total number of TEs called and found them to be positively correlated (Figure 6a) with R^2^ = 0.88 and *p* < 0.001. Looking at the totals of individual TE families yielded similar results with R^2^ = 0.87 for Alus, R^2^ = 0.90 for L1s, R^2^ = 0.92 for SVAs and R^2^ = 0.90 for HERVs (all *p* < 0.001) (Figure 6b).

**Figure 6:**
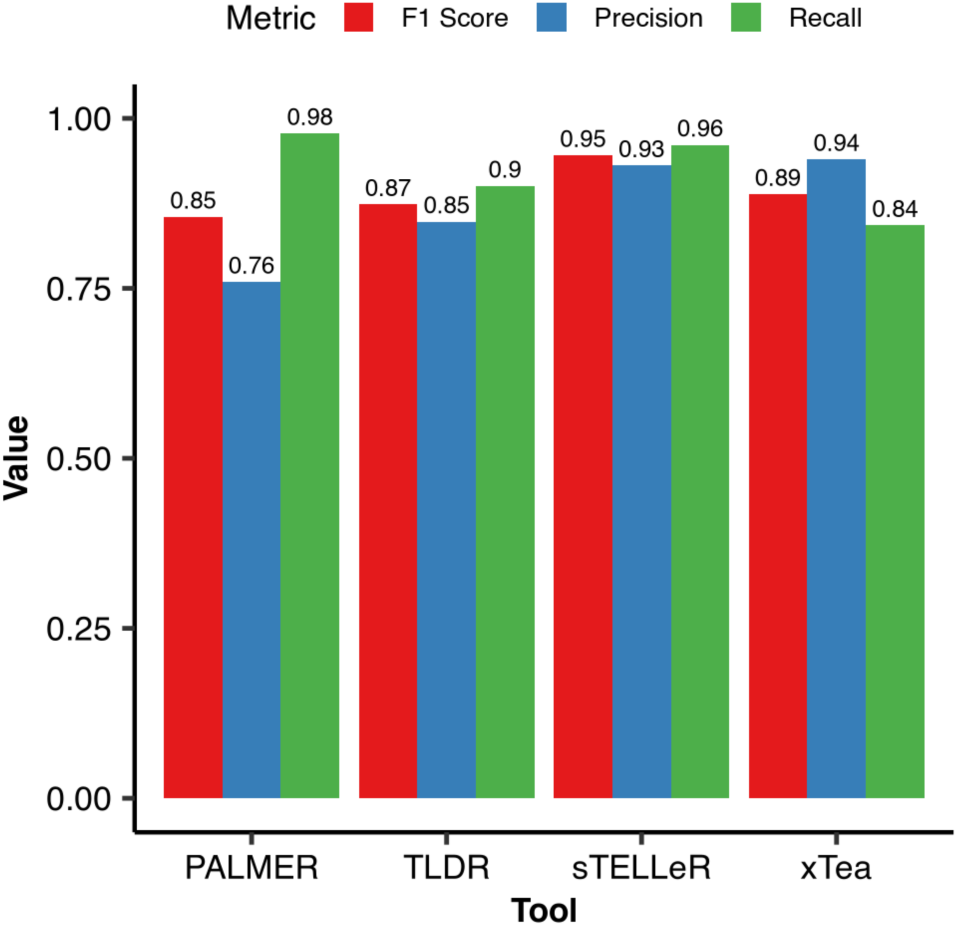
Performance metrics for GIAB data for the refined truth set. a. Precision, Recall and F1-scores for PALMER, TLDR, sTELLeR and xTea for the GIAB data using calls that are called by 3 out of 4 tools and are in the high confidence SV list as truth set.

As coverage is associated with the samples being singleplex or multiplex, we can also observe this trend when we look at the total number of TEs called for each sample (Figure 7b). There are expected to be over a million Alu copies in the genome (this includes reference insertions), 500,000 L1 copies, 3000 SVA copies and very few active HERV copies[25]. This trend can be observed in the total number of TEs called in different families as well, with mean Alu calls by the tools is 57085, L1 is 9005, SVA is 3056 and HERV is 1063. However, these numbers are highly influenced by the over-calling of PALMER as can be seen in Figure 5a. In PALMER’s documentation, the authors suggest using high confidence reads (HCR) ≥ 1 and potential supporting reads (PSR) ≥ 10% of average coverage. In the multiplex samples, this equates to just 1 PSR which we thought was not enough support to call a TE, therefore we have opted for ≥ 2 as the PSR threshold. Even so, we have observed a highly inflated number of TEs called by PALMER. More surprisingly, amongst the calls from all 14 samples, we had no HCR support for any of the TEs, so we ignored that threshold. TLDR is the most stringent caller of the four as it calls the fewest in all TE families except SVA, in which it is very close to xTea’s 728 with 747. When we split TE family calls into samples, we can confirm that lower coverage that comes with being multiplex samples does not change the Alu > L1 > SVA > HERV distribution of number of calls (Figure 7c).

**Figure 7:**
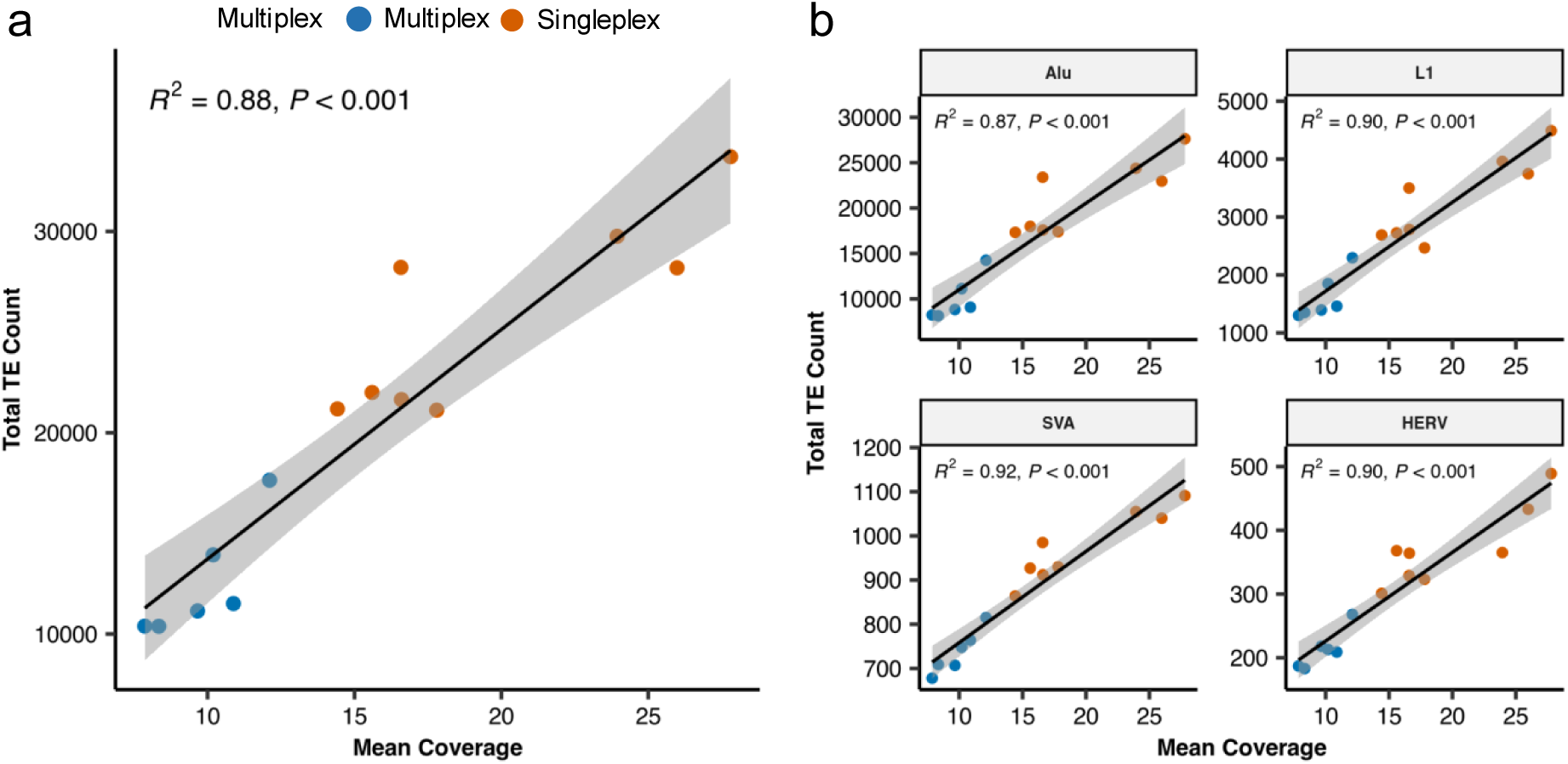
Correlation of total number of TEs detected vs. mean coverage. a. for all TEs. b. for different TE families.

To assess the concordance and divergence among TE detection tools, we compared the insertion calls made by PALMER, sTELLeR, TLDR, and xTea across the full dataset (Figure 8). Notably, PALMER uniquely reported 158,167 insertions, comprising 56.3% of all TE calls, suggesting a highly permissive calling strategy that may prioritize recall at the potential cost of precision. In contrast, xTea reported only 72 insertions not identified by any other tool (0.0%), reflecting a more conservative approach likely aimed at minimizing false positives. A total of 69,209 insertions (24.6%) were identified by all four tools, representing a core set of high-confidence insertions. Intermediate overlaps—such as 25,510 insertions shared by PALMER and sTELLeR, and 13,006 shared by PALMER and xTea—further indicate partial concordance between certain tools (Figure 8a). TLDR exhibited very few tool-exclusive calls, suggesting a preference for insertions supported by multiple sources.

**Figure 8:**
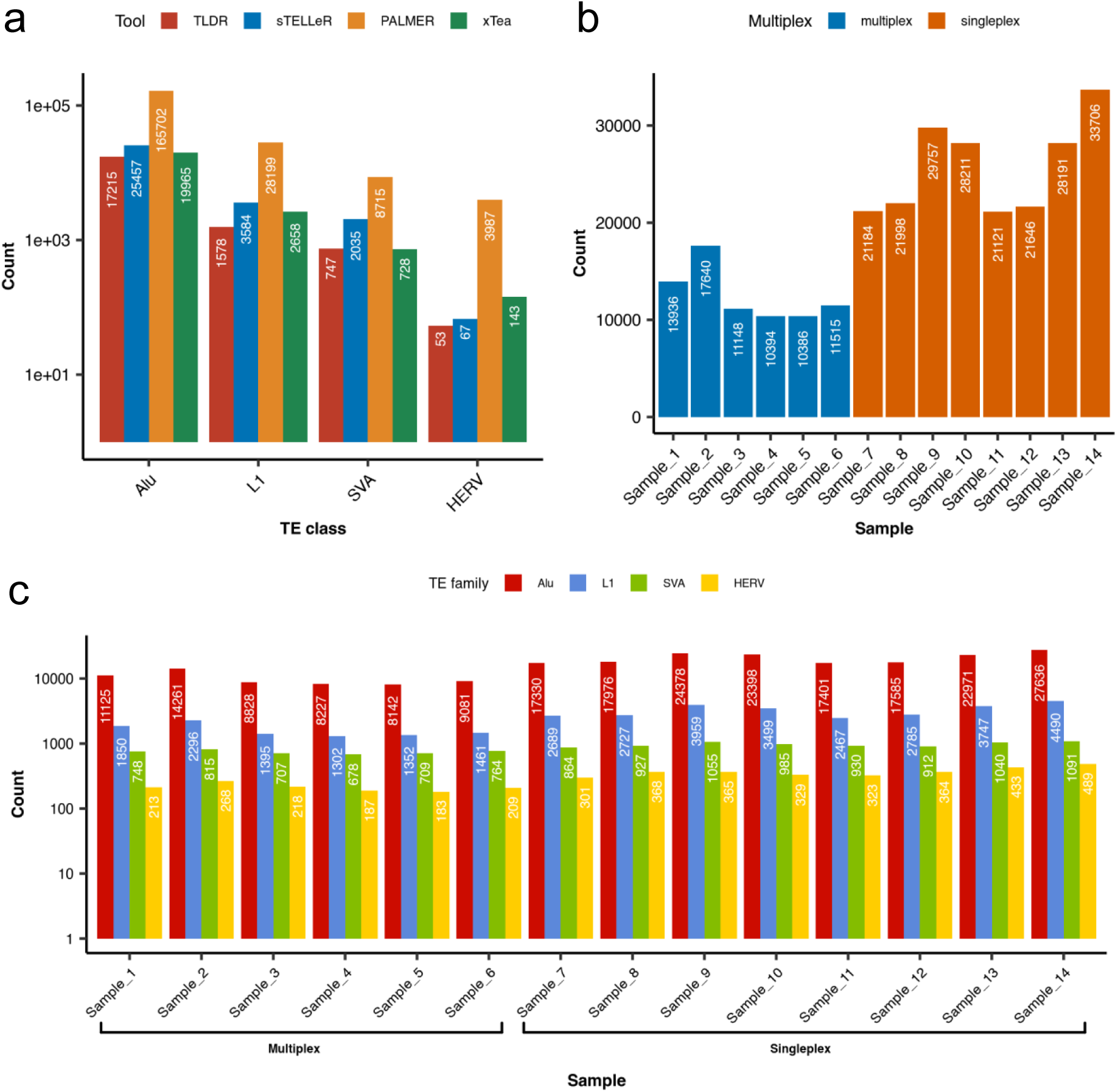
Distribution of detected TEs. a. for different TE families with different tools. b. of total number of TE calls by singleplex/multiplex c. by singleplex vs multiplex samples for different TE families.

These results collectively highlight the trade-offs between stringency and recall across TE detection algorithms and underscore the importance of integrating multiple tools or applying post hoc validation to achieve a reliable call set. To further contextualize the level of consensus among tools, we examined the fraction of each tool’s total calls that overlap with the 69,209 insertions identified by all four tools. TLDR exhibited the highest concordance, with 87.6% of its calls supported by all tools, followed by xTea (78.1%) and sTELLeR (59.8%), suggesting a high degree of precision and agreement with other methods. In contrast, only 24.9% of PALMER’s total calls were shared by all tools, reinforcing its tendency to call a large number of tool-specific insertions and potentially reflecting a more permissive or less stringent calling strategy. Across all TE families, PALMER consistently contributed the lowest fraction of insertions shared by all tools (e.g., 5.9% in SVA (Figure 8d), 16.3% in L1(Figure 8c)), while xTea exhibited the highest agreement in Alu (91.6% (Figure 6a)) and L1 (81.1%), supporting its conservative and high-precision profile. Notably, HERV insertions have better consensus across all tools, with sTELLeR and TLDR having zero calls not identified by any other tool and xTea having only two, possibly highlighting the inherent advantage of using LRS in reliably detecting endogenous retroviruses (Figure 8e).

There are multiple TE database search tools which have valuable information about transposable elements. RepeatMasker[26] is one of them, having information about interspersed repeats and low complexity DNA sequences. Telescope[27] and ERVmap[28] are specialized databases that have information on HERVs. We also compared our results with these databases to account for the possibility of overlap with known TEs. Although the tools we benchmark focus on non-reference TE detection, not all non-reference TEs represent novel insertions; some may correspond to variants of reference TEs. In general, there is a high overlap with RepeatMasker for all tools, meaning there is quite a fair percentage of called TEs that are known. As expected, there is little to no overlap between detected Alu, L1 and SVAs and the HERV specific databases. The little overlap (1.7% at most in sTELLeR between SVA and ERVmap) that these calls have with the databases could be the result of wrong annotation by the tools or the databases themselves as these databases were also curated using short read sequencing techniques, which as we mentioned before could prove inaccurate when mapping low complexity regions or longer insertions. Surprisingly, there is at most 26.9% with sTELLeR overlap between called TEs and HERV specific databases. TLDR’s overlap is also 26.4% in comparison to 11.2% in xTea and 6.2% in PALMER (Figure 9). PALMER results can also be attributed again to its lenient filtering and large number of calls.

**Figure 9:**
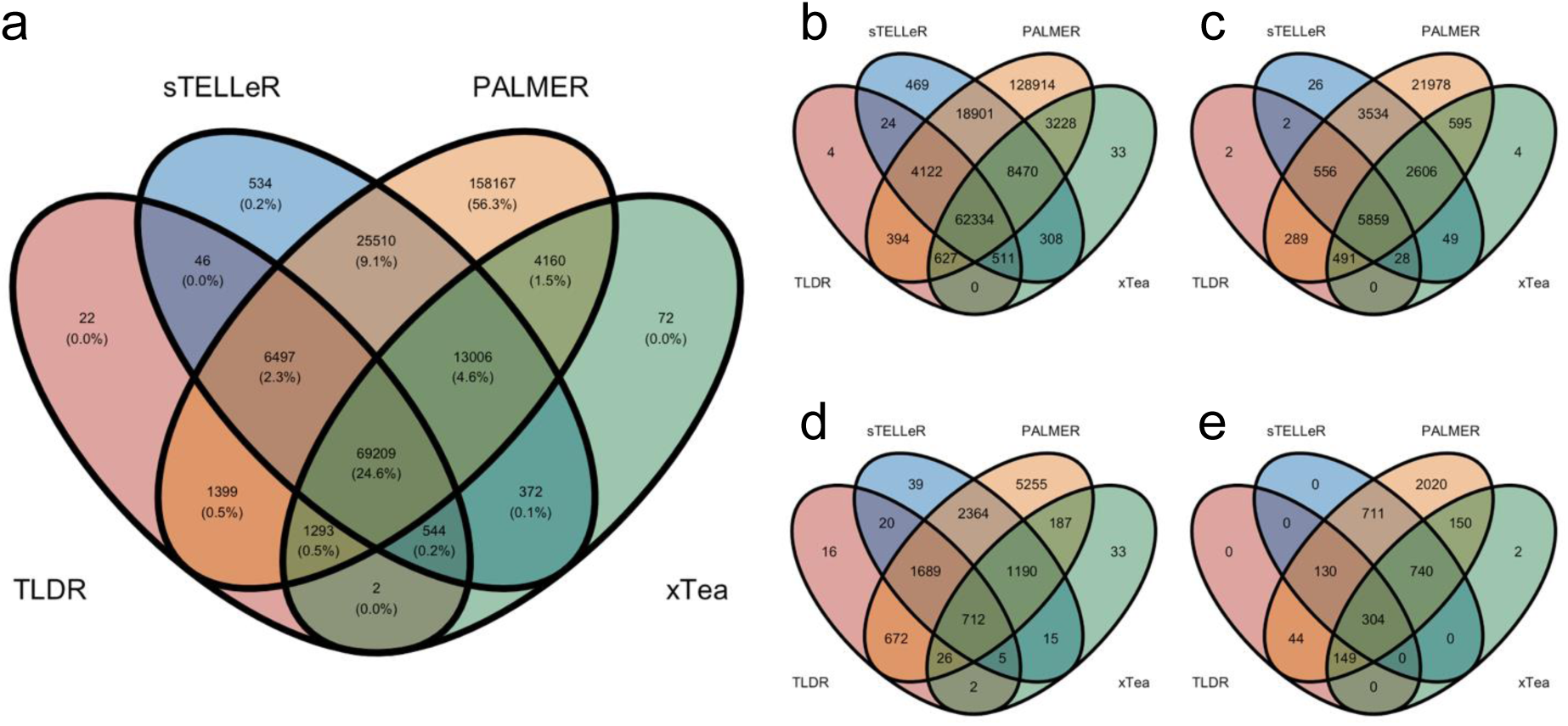
Venn diagrams showing the overlap of detected TEs between the four tools. a. For all TEs. b. for Alu. c. for L1. d. for SVA. e. for HERV.

For these 14 samples, we also had SRS data[24] and SRS TE calls. The Alu, L1 and SVA calls from SRS were made by MELT[29], whereas HERV calls were made by HERV-specific detection tools STEAK[30] and RetroSnake[31,32]. We utilized these TE calls to compare SRS and LRS results and checked the percentage of long-read calls that are supported by short-read calls and vice-versa, short-read calls that are supported by long-read calls. Overall, about 40-50% of the calls by LRS tools are also detected with SRS (Figure 10a). The permissiveness of PALMER plays a role here again, causing at best ∼25% coverage by SRS in HERVs, dropping dramatically in other TE families. xTea appears to have the best SRS coverage, especially in SVA where almost 75% of the calls are supported by SRS. On the other hand, when we look at short-read calls supported by the 4 tools, we see a better overlap, especially for PALMER where 81% of Alu and 85% of SVA calls from SRS are detected (Figure 10b). This indicates yet again that while PALMER detects TEs accurately, the sheer amount of calls it makes reduces its overall reliability. Considering LRS calls as higher confidence TE calls with respect to SRS, this experiment suggests that although short-read TE calls might be high confidence, still 50% or more TEs remain undetected by SRS. Interestingly, we see extremely low overlap in HERVs which may be caused by the difference in tools used as well.

**Figure 10:**
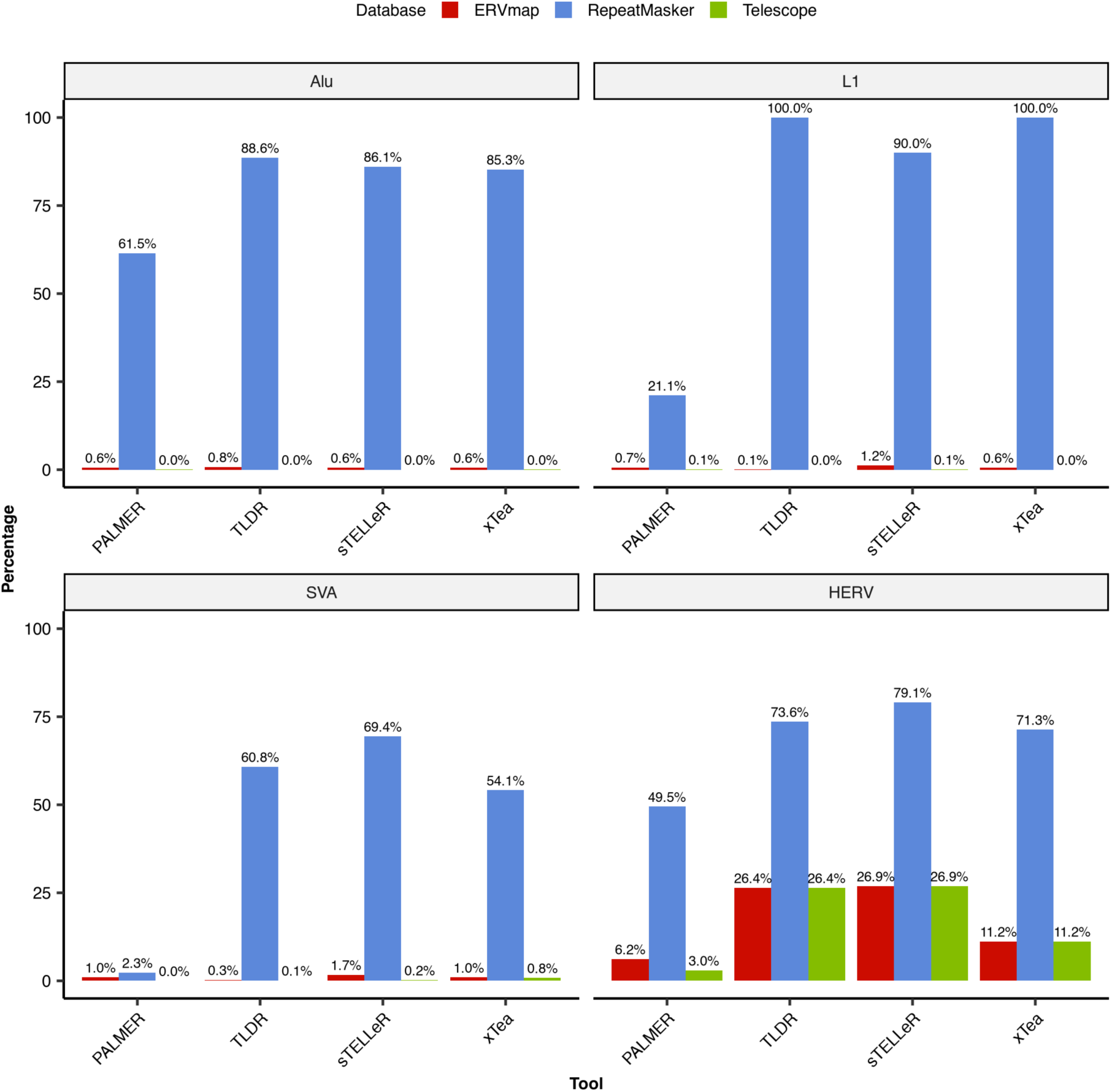
Overlap percentages of Brain Bank TE calls with ERVmap, Repeatmasker and Telescope databases.

We also used gnomAD-SV 4.0[33] to check the population allele frequencies (AF) of the detected TEs. Our rationale behind this was that real TEs would tend to have higher population AF in the gnomAD-SV database comparing with calling mistakes[34,35]. For this, we split the AFs into three groups: Low: AF < 0.01, Medium: 0.01 – 0.1 and High: ≥ 0.1. Looking at the distribution of these groups of AFs in the four tools shows that almost three quarters of TEs detected by xTea belong to the High AF group, followed by sTELLeR, TLDR and further low PALMER (Figure 11). These results are in line with the previous analyses where most calls made by PALMER appeared to be artifacts. It also re-establishes xTea and sTELLeR’s statuses as the most stringent tools, once again shown that their calls are more likely to be real TEs. An interesting observation is that TLDR has by far the most Medium AF TEs with more than twice of its closest rival tool. This AF range often encompasses population-specific or recently arisen polymorphisms, suggesting TLDR may be more sensitive to such variants. While this could indicate improved detection of biologically relevant insertions, it may also reflect differences in the handling of complex genomic regions, warranting further targeted validation.

**Figure 11:**
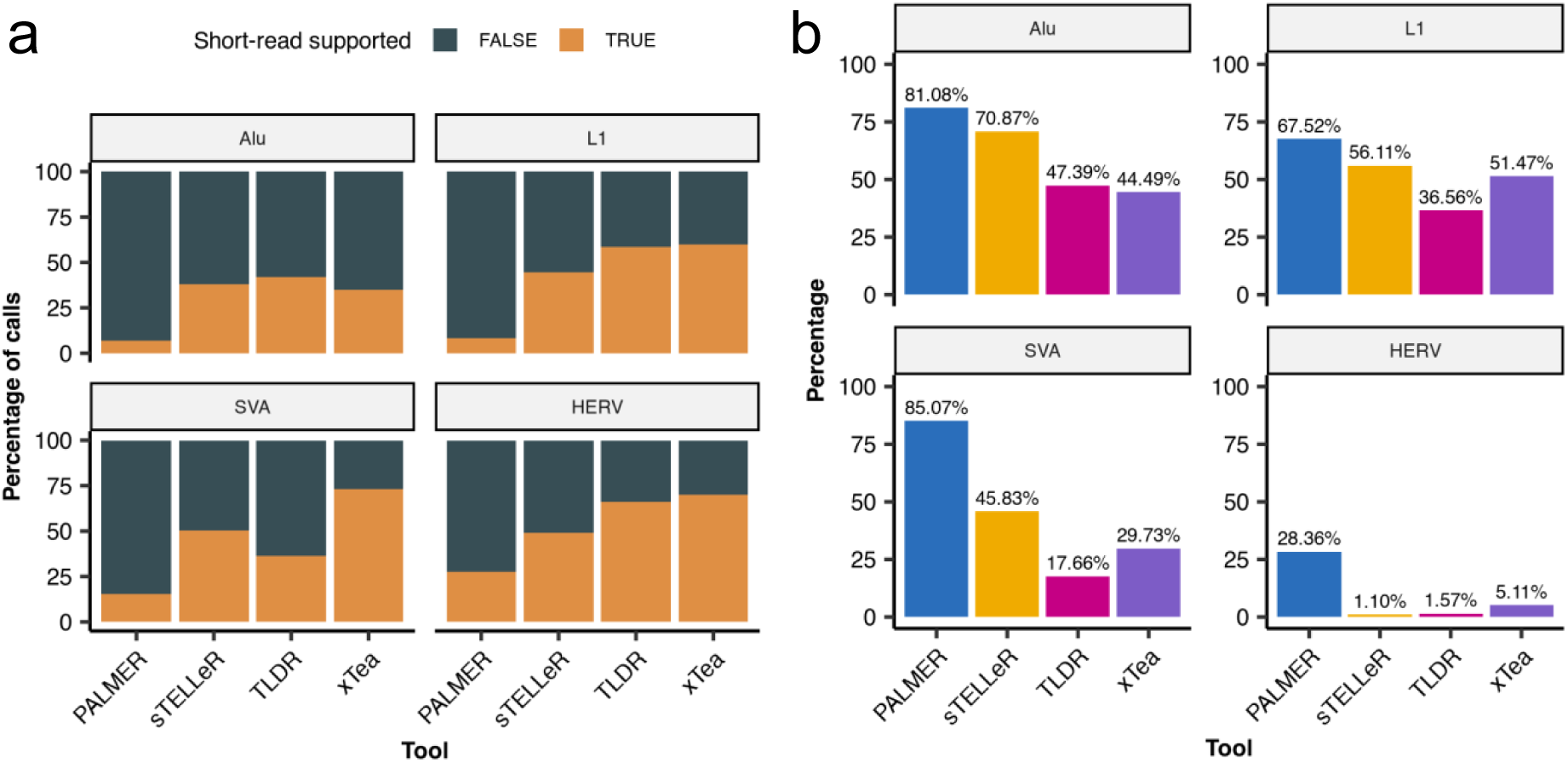
Short read support for the long read TE calls. a. Percentage of TEs detected by the benchmarked tools that are common with the short read calls. b. Percentage of long read support for the TEs detected by short read sequencing.

When we look at the frequencies of the different genotypes of the TEs called by the four tools, we see that sTELLeR and xTea have relatively low frequencies of homozygous reference (FREQ_HOMREF) compared to the other two tools. PALMER’s calls are predominantly homozygous reference, pointing to a bias towards reference TEs instead of non-reference TEs. In comparison, sTELLeR and xTea have the highest homozygous alternative genotype frequencies as well as the heterozygous genotype frequencies, leading us to believe more of these two tools’ TE calls are valid non-reference TEs (Figure 12).

**Figure 12:**
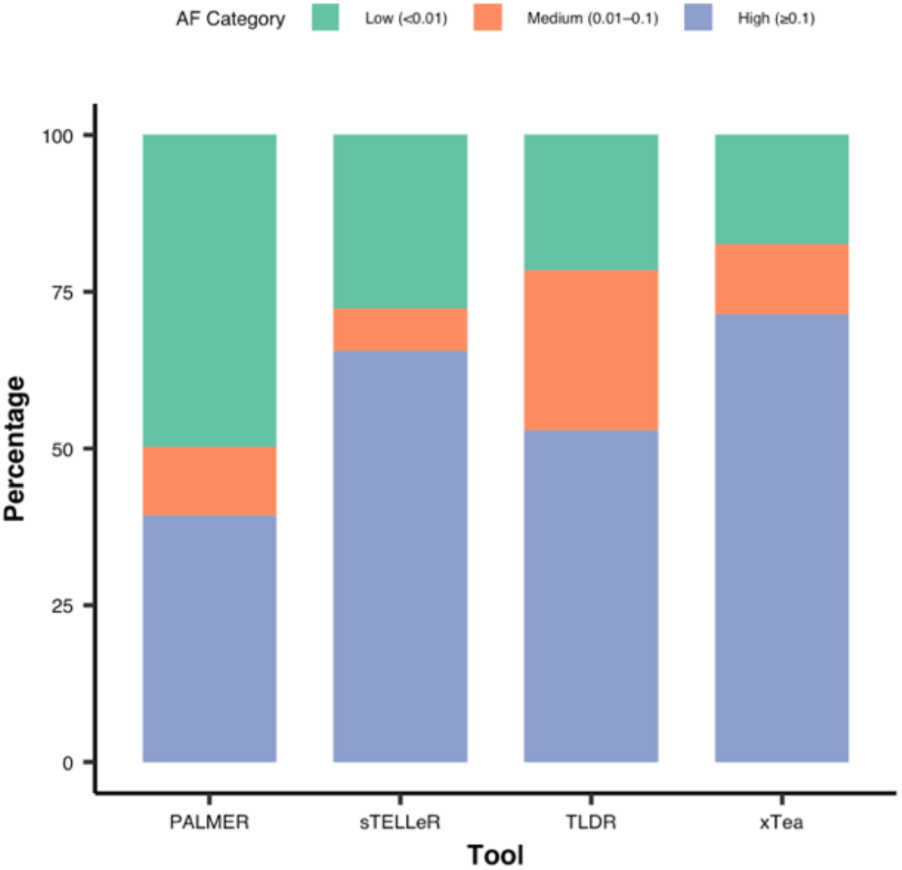
gnomAD allele frequency group percentages (Low: AF < 0.01, Medium: 0.01 – 0.1, High: ≥ 0.1).

**Figure 13:**
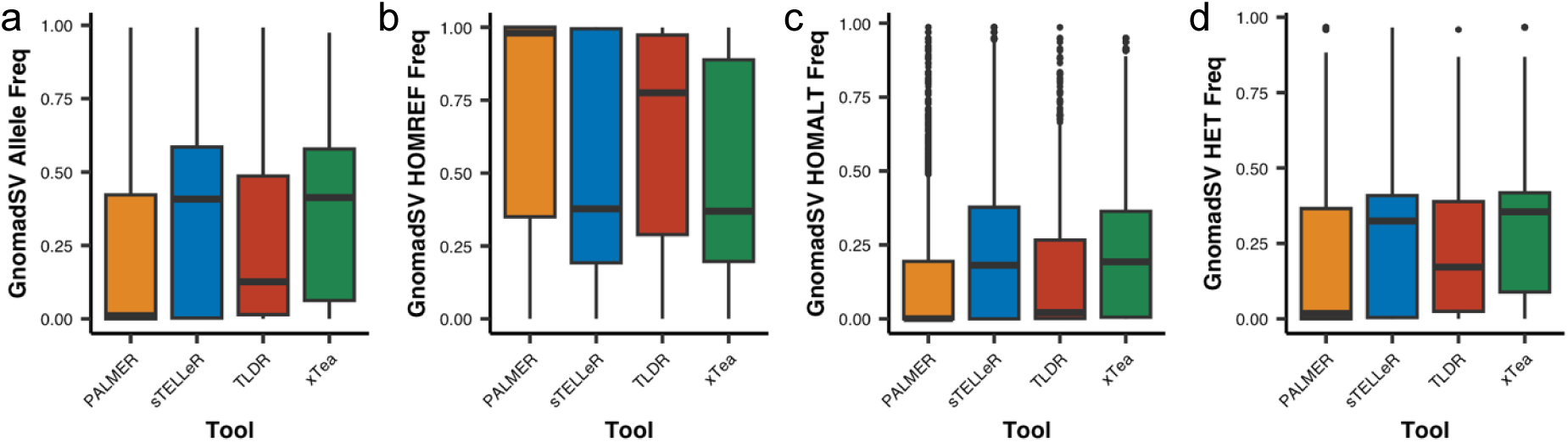
gnomAD allele and genotype frequency distributions. a. Allele frequencies of TEs detected. b. Homozygous reference genotypes frequencies. c. Homozygous alternative genotype frequencies. d. Heterozygous genotype frequencies.

## Discussion

Benchmarking of transposable element (TE) detection tools remains essential for informing tool selection and optimizing variant discovery in large-scale genomics projects. Using the well-characterized GIAB HG002 sample as a reference, along with simulated data and 14 LND BB samples, we systematically compared the performance of four long-read TE detection tools — PALMER, TLDR, sTELLeR, and xTea. Our evaluation highlights distinct trade-offs in precision, recall, and population-level support, underscoring that no single tool outperforms across all criteria.

### Performance metrics reveal divergent calling strategies

Across F_1_-score, precision, and recall, the four tools demonstrated complementary strengths. TLDR and xTea showed the highest precision among single callers, albeit at the cost of reduced recall, consistent with more stringent filtering of candidate insertions. PALMER achieved the highest recall, recovering the largest fraction of known TE insertions, but with a corresponding reduction in precision. sTELLeR displayed balanced intermediate performance, capturing a wide range of events while maintaining moderate precision. Notably, combining calls from multiple tools — particularly intersections of three or more callers — improved precision to near 100%, suggesting that consensus-based approaches may be an effective strategy when minimizing false positives is the primary objective. However, TLDR and xTea, despite their high single-tool precision, contributed fewer events to such high-precision intersections, reflecting their conservative calling thresholds.

### Population allele frequency profiles differentiate tool outputs

Integration with gnomAD-SV data provided additional insight into the population prevalence of detected TEs. Grouping calls by allele frequency (AF) revealed that TLDR was enriched for insertions in the medium AF range (0.01–0.1), nearly doubling the proportion of such events compared to other tools. This may reflect TLDR’s tendency to preferentially detect variants with moderate population support — potentially representing recurrent but not ubiquitous insertions. In contrast, PALMER was skewed toward low AF (<0.01) events, consistent with its high recall and capacity to recover rare or potentially sample-specific insertions. In interpreting this result it is important to highlight that sequencing mistakes are more likely to result in incorrectly called rare variants[36–38] which means PALMER’s higher recall might come at the expense of its precision. sTELLeR and xTea detected a greater proportion of high AF (≥0.1) events, aligning with their capture of well-supported, common insertions.

### Genotype distribution supports tool-specific biases

Examining gnomAD-SV genotype frequencies further supported these patterns. sTELLeR and xTea had lower median homozygous reference (HOMREF) frequencies compared to PALMER and TLDR, indicating that their calls are present in a larger fraction of individuals in the population. PALMER’s low median homozygous alternate (HOMALT) frequency points to its enrichment for rare alleles, while xTea’s higher HOMALT values suggest detection of common, often fixed insertions. Heterozygous (HET) frequencies were broadly similar across tools, but PALMER’s slightly lower values again support a bias toward rarer events.

### Comparison with short-read calls and computational performance

Our experiments using 14 ONT samples with varying coverage levels highlight how sequencing depth, tool choice, and platform comparisons strongly influence TE detection outcomes. We observed a clear positive correlation between coverage and the number of TEs called across all families, consistent with expectations given the repetitive and low-mappability nature of these elements. While singleplex samples with ∼20× coverage yielded more insertions than multiplexed samples at ∼10×, the relative abundance across TE families (Alu > L1 > SVA > HERV) remained stable regardless of coverage, indicating that depth primarily affects sensitivity rather than biasing family distribution. Comparison with short-read data further revealed that only ∼40–50% of long-read calls were supported by SRS, underscoring the limitations of short-read methods in repetitive regions. Tool performance varied substantially: xTea demonstrated the best concordance with SRS in SVA calls (∼75% overlap), whereas PALMER’s permissive thresholds produced inflated totals and low validation rates, particularly for HERVs. Interestingly, PALMER still recovered the majority of Alu and SVA insertions detected by SRS, suggesting high sensitivity but limited specificity. From a computational standpoint, all tools are directly influenced by coverage/input file size, with increased coverage resulting in increased resource requirements and runtimes. However, computational performance of a tool is neither an indicator of tool performance nor the number of TEs a tool might call. While sTELLeR overall calls the 2^nd^ highest number of TEs in the 14 LND BB samples, it is multiples faster and less resource intensive than its competitors. Taken together, these findings indicate that analysis design must balance sequencing depth, computational resources, and tool selection: higher coverage improves sensitivity, but stringent callers or consensus approaches may be preferable when the goal is robust, reproducible TE discovery.

### Practical implications for tool choice

From a practical perspective, these differences in detection profiles have important implications. Studies aiming to catalogue rare or novel TE insertions — for example, in rare disease discovery or population-specific variant analysis — may benefit from PALMER’s broad recall and TLDR’s capacity to capture moderately frequent alleles. Conversely, projects prioritizing confident identification of common insertions, such as population reference panel construction, may find sTELLeR and xTea outputs more suitable. Given the observed benefits of multi-caller consensus, hybrid approaches incorporating two or more tools could leverage these complementary strengths while mitigating individual weaknesses.

### Limitations and future directions

Our benchmark relied on a single reference sample (HG002) with high-quality truth annotations, which may not capture the full complexity of TE diversity across populations. Long-read sequencing depth, platform-specific error profiles, and parameter tuning were not exhaustively explored and could influence performance rankings. Moreover, gnomAD-SV provides population-level support primarily from short-read data, which may underrepresent certain TE types or genomic contexts. Expanding benchmarks to multiple ancestries, sequencing platforms, and TE classes will be critical to generalizing these findings. Two more recently developed tools LOCATE and TrEMOLO were also not included, one for being under review, the other due to integration issues at the time of writing this manuscript.

## Conclusion

This comprehensive evaluation highlights that tool choice for TE detection involves clear trade-offs between recall, precision, and population support. While PALMER and TLDR are better suited for detecting rare insertions, sTELLeR and xTea excel at identifying common, well-supported events. Consensus-based approaches can maximize precision, though at the cost of reduced recall. Our results provide a practical framework for an informed selection of TE detection strategies tailored to specific research goals and underscore the value of integrating population allele frequency data to contextualize variant calls.

## Methods

### Code availability

Scripts used to run tools and analyze results are available upon request. We have omitted parameter tuning and used the same TE reference (https://github.com/kristinebilgrav/sTELLeR_supplementary/blob/main/fasta/TEseque nces_SVA_HERV_ALU_L1.fasta) for all tools that allowed for custom TE reference. For PALMER where it is necessary to run separately for each TE family, this fasta was split into different files containing the separate TE family sequences.

### Tool selection

We have used all available tools (PALMER, TLDR, sTELLeR, xTea) that specialize on non-reference TE detection using long reads to our knowledge. There are two other tools LOCATE and TrEMOLO which have been recently developed in 2025. We have excluded them from our benchmark as LOCATE is still in preprint and is being revised and we could not integrate TrEMOLO to our workflow King’s College London’s HPC CREATE[39] due to repetitive errors while running the tool.

### Data

#### Simulated data

We have used the GRCh38 reference genome with the alternate locus scaffold omitted (`GCA_000001405.15_GRCh38_no_alt_analysis_set.fna)` to insert random TEs and create the simulated reads using pbsim3. We have used the QSHMM-ONT-HQ model constructed from ONT reads. minimap2[40] was used to align the simulated reads to the reference.

#### Real data

We have used the publicly available GIAB v0.6 Tier 1 benchmark set for the HG002 sample as part of our benchmarking of real world data, thanks to it being extensively analysed and having carefully curated call sets. The benchmark set was obtained from https://ftp-trace.ncbi.nlm.nih.gov/ReferenceSamples/giab/data/AshkenazimTrio/analysis/NIST_SVs_Integration_v0.6/ and the PALMER L1 calls for the same sample from https://ftp-trace.ncbi.nlm.nih.gov/ReferenceSamples/giab/data/AshkenazimTrio/analysis/PacBio_PALMER_11242017/.

In addition, we used 14 ONT samples from the LND BB for which the raw data requests can be addressed to Prof. Alfredo Iacoangeli as the data is part of an ongoing study. For Oxford Nanopore Technologies (ONT) sequencing, genomic DNA was extracted using the Monarch® Spin gDNA Extraction Kit (New England Biolabs, Hitchin, UK) and libraries were prepared with the SQK-LSK114 Ligation Sequencing Kit (Oxford Nanopore Technologies, Oxford, UK), following the manufacturers’ protocols. Library concentrations were measured with a Qubit fluorometer (Thermo Fisher Scientific). For multiplexed sequencing, samples were barcoded with the SQK-NBD114.24 Native Barcoding Kit (Oxford Nanopore Technologies). Sequencing was performed on PromethION 24 instruments using R10.4.1 flow cells, with two samples pooled per flow cell for multiplexed runs, and single samples run per flow cell for singleplexed experiments.

EPI2ME wf-basecalling v1.4.5 pipeline[41] was used with the Dorado model dna_r10.4.1_e8.2_400bps_sup_v5.0.0 to base-call the raw signal data. Reads were then aligned to the GRCh38 reference genome using the EPI2ME wf-human-variation v2.6.0 pipeline[42]. Performance analysis, plots and figures were done in R programming language (v4.4.2)[43], using R Studio (v 2025.09.0+387)[44].

Short-read samples were generated using Illumina’s FastTrack service (Illumina, San Diego, CA, US) using the Illumina HiSeq 2000 platform. Isaac was used to align the 100 bp reads generated by PCR-free library to hg19 with approximately 35x coverage per sample.

HERVs from SRS data were called using RetroSnake, which is a Snakemake workflow for RetroSeq, using default parameters and both *discover* and *call* options active. Resulting calls were filtered using the following approach: For *fl* of 8, *gq* has to be 10 or more; for *fl* of 7 *gq* has to be 20 or more and for *fl* of 6, *gq* needs to be as high as 29. All predictions with *fl* lower than 6 are rejected[31]. Alu, L1 and SVAs were called using MELT (v2.2.2) with default parameters. TEs with a minimum size of 50bases were considered for the downstream analysis. Both RetroSnake and MELT results were later lifted over to GRCh38 using UCSC Genome Browser’s liftOver tool[45] to match the results of the LRS TE calls.

#### Execution of detection tools and computational requirements

A Nextflow[46] workflow was created to run all tools for the Brain Bank samples with the parameters “-with-report” and “-with-trace” to keep track of all resource usage and runtime information. xTea and TLDR are the only tools capable of multithreading, and they were run with 8 threads, while sTELLeR and PALMER do not have thread option so were run with single threads.

#### Performance metrics

We have used various performance metrics to compare the abilities of TE callers. We measured Precision, to measure accuracy of positive predictions, Recall/Sensitivity, to measure a tool’s ability to find all positive instances, and finally F1-score, the harmonic mean of precision and recall, giving a good overall metric to assess performance.

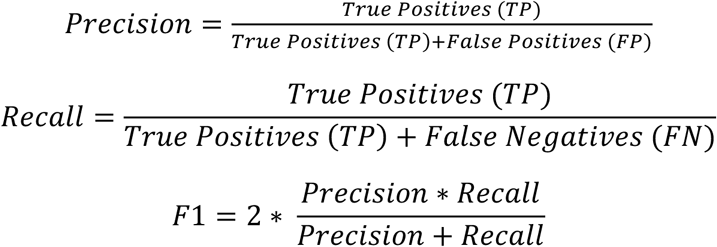

For the correlation of coverage vs. number of TEs called, we fitted a linear model, using the *stat_poly_eq* and *stat_poly_line* functions from the ggpmisc (v0.6.1)[47] package in R.

## Acknowledgments

Samples used in this research were in part obtained from the UK National DNA Bank for MND Research, funded by the MND Association and the Wellcome Trust. We thank people with MND and their families for their contribution to the UK National DNA Bank for MND Research. The authors acknowledge use of the King’s Computational Research, Engineering and Technology Environment (CREATE) (https://create.kcl.ac.uk, accessed on 5 May 2025), which is delivered in partnership with the National Institute for Health and Care Research (NIHR) Biomedical Research Centres at South London and Maudsley and Guy’s and St. Thomas’ NHS Foundation Trusts and part-funded by capital equipment grants from the Maudsley Charity (award 980) and Guy’s and St. Thomas’ Charity (TR130505). We also acknowledge Health Data Research UK, which is funded by the UK Medical Research Council, Engineering and Physical Sciences Research Council, Economic and Social Research Council, Department of Health and Social Care (UK), Chief Scientist Office of the Scottish Government Health and Social Care Directorates, Health and Social Care Research and Development Division (Welsh Government), Public Health Agency (Northern Ireland), British Heart Foundation and Wellcome Trust. The London Neurodegenerative Diseases Brain Bank at KCL has received funding from the MRC and through the Brains for Dementia Research project (jointly funded by Alzheimer’s Society and Alzheimer’s Research UK.

