## Supplementary figures and images for "Systematic comparative benchmarking of computational methods for the detection of transposable elements in long-read sequencing data"

### Supplementary Figure 1

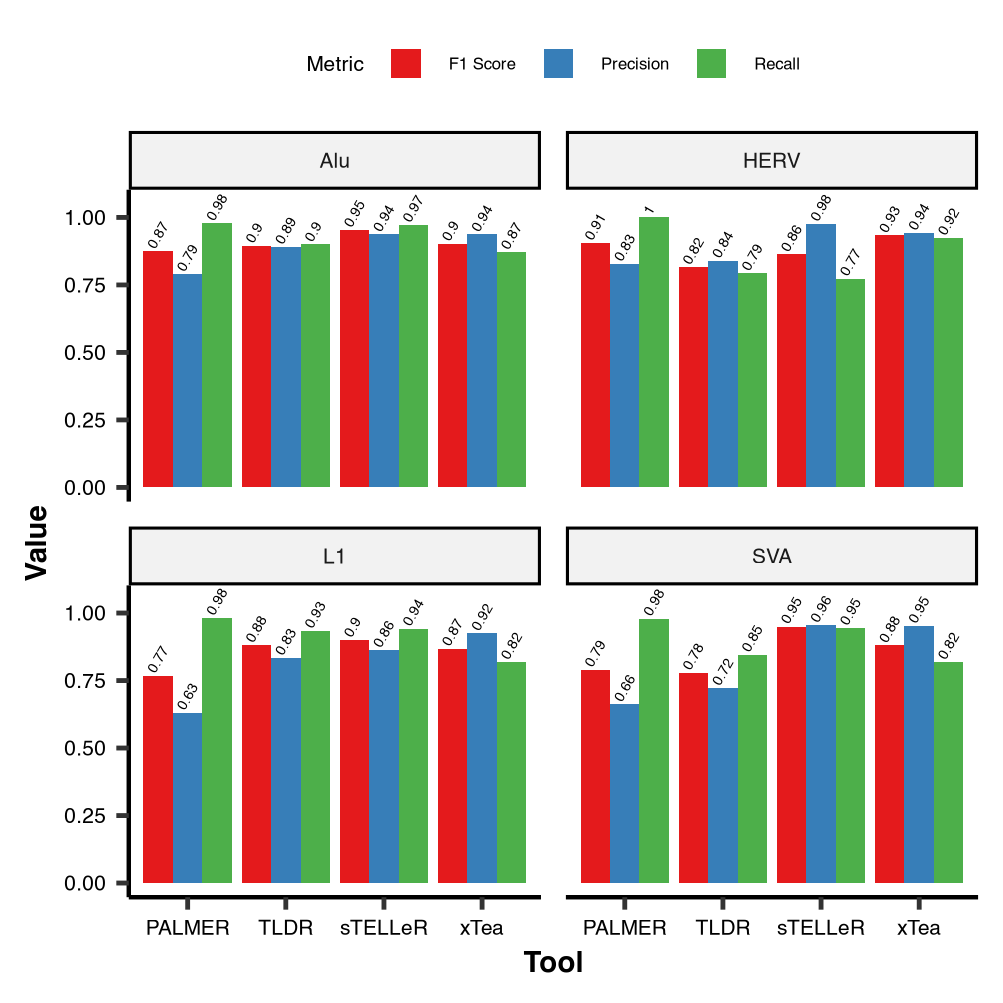

### Supplementary Figure 2

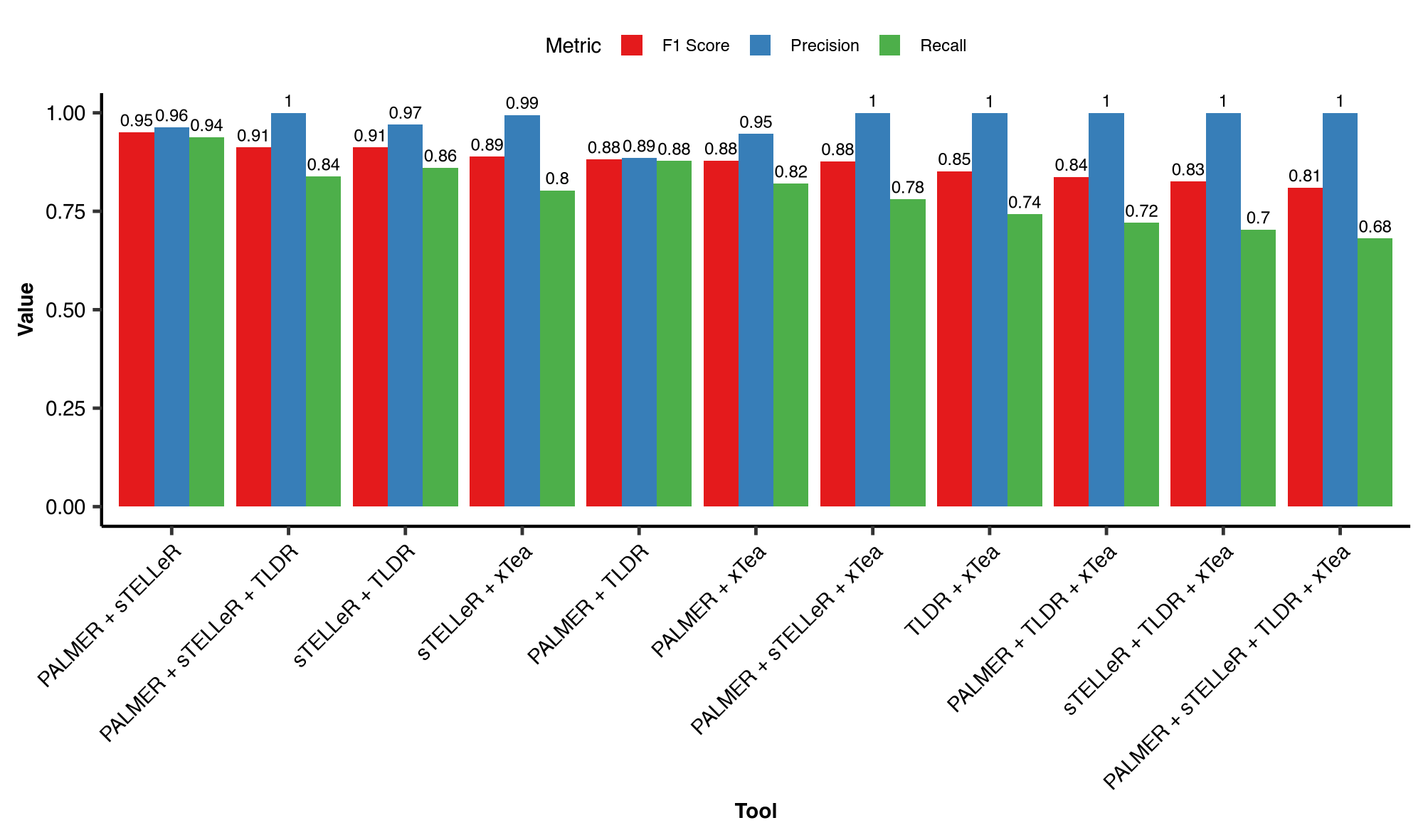
